# Evidence accumulation determines conscious access

**DOI:** 10.1101/2020.07.10.196659

**Authors:** Michael Pereira, Pierre Megevand, Mi Xue Tan, Wenwen Chang, Shuo Wang, Ali Rezai, Margitta Seeck, Marco Corniola, Shahan Momjian, Fosco Bernasconi, Olaf Blanke, Nathan Faivre

**Author notes:** equal contribution (Olaf Blanke, Nathan Faivre). Corresponding author: Nathan Faivre, Laboratoire de Psychologie et Neurocognition, CNRS UMR 5105, UGA BSHM, 1251 Avenue Centrale, 38058 Grenoble Cedex 9. Author Contributions: MP, MT, and NF developed the study concept and contributed to the study design. PM and MS developed the clinical trial that generated the microelectrode array data. Data collection and data analysis were performed by MP, PM, FB, MT, WC, and NF. MP performed modelling work. MC and SM performed surgery. MP and NF drafted the paper; all authors provided critical revisions and approved the final version of the paper for submission.

## Abstract

A fundamental scientific question concerns the neuronal basis of perceptual consciousness, which encompasses the perceptual experience and reflexive monitoring associated with a sensory event. Although recent human studies identified individual neurons reflecting stimulus visibility, their functional role for perceptual consciousness remains unknown. Here, we provide neuronal and computational evidence indicating that perceptual and reflexive consciousness are governed by an all-or-none process involving accumulation of perceptual evidence. We recorded single-neuron activity in a participant with a microelectrode implant in the posterior parietal cortex, considered a substrate for evidence accumulation, while he detected vibrotactile stimuli around detection threshold and provided confidence estimates. We found that detected stimuli elicited firing rate patterns resembling evidence accumulation during decision-making, irrespective of response effectors. Similar neurons encoded the intensity of task-irrelevant stimuli, suggesting their role for consciousness per se, irrespective of report. We generalized these findings in healthy volunteers using electroencephalography and reproduced their behavioral and neural responses with a computational model. This model considered stimulus detection if accumulated evidence reached a bound, and confidence as the distance between maximal evidence and that bound. Applying this mechanism to our neuronal data, we were able to decode single-trial confidence ratings both for detected and undetected stimuli. Our results show that the specific gradual changes in neuronal dynamics during evidence accumulation govern perceptual consciousness and reflexive monitoring in humans.

---

The processing of sensory signals by the human brain gives rise to two interrelated phenomena: perceptual consciousness, defined as the subjective experience associated with a sensory event (Chalmers, 1995; Nagel, 1974; Block 2011), and perceptual monitoring, defined as the capacity to introspect and reflect upon the subjective experience associated with a sensory event (Flavell, 1979; Koriat, 2006; Fleming, Dolan, and Frith 2012). The main strategy employed to study conscious processing consists in relating first-order subjective reports to neural activity to identify the minimal set of neuronal events and mechanisms sufficient for a specific conscious percept (i.e., neural correlates of consciousness or NCCs : Koch et al., 2016). To identify NCCs, most experimental paradigms have adopted a contrastive approach, whereby distinct phenomenal experiences induced by constant sensory stimulation are compared (Baars, 1998). One of the simplest contrasts is obtained when stimuli are presented at low intensity or embedded in noise so that only a certain proportion of them is detected (Dehaene et al., 2006). A comparison of neural activity elicited by detected and missed stimuli allows distinguishing the neural correlates of conscious vs. unconscious sensory processing, and therefore identifying NCCs given that specific confounds are ruled out (Aru et al., 2012). However, although rare investigations in humans have described single neurons in the temporal lobe encoding stimulus detection (Quiroga et al., 2008; Reber et al., 2017), the mechanistic role of neuronal activity for perceptual consciousness remains unknown. One prominent theory of consciousness, the global neuronal workspace, proposes that a stimulus is consciously perceived when its corresponding neural activity is globally broadcasted across the cortex (Mashour, et al., 2020). This theory assumes that this global broadcast is triggered when an (unconscious) evidence accumulation process reaches a threshold (Dehaene et al., 2014; Dehaene, 2009; Shadlen 2011), similar to the physiological processes underlying decision-making (Bollimunta & Ditterich, 2012; Katz et al., 2016; Roitman & Shadlen, 2002; Zhou & Freedman, 2019). Although various neuroimaging studies have interpreted increases in neural activity elicited by detected stimuli (versus missed stimuli) as evidence accumulation (Salti et al., 2015; Tagliabue et al., 2019; Wyart & Tallon-Baudry, 2009), little empirical evidence supports an evidence accumulation account of perceptual consciousness, especially at the single neuron level.

Besides perceptual consciousness, the main strategy to study perceptual monitoring consists in assessing how second-order reports like confidence judgments co-vary with the accuracy of a given perceptual task (first-order reports; Fleming & Lau, 2014). As most studies investigating perceptual monitoring rely on first-order discrimination tasks with stimuli that are always detected, less is known regarding how the brain monitors the presence or absence of subjective experience (Li et al., 2014; Mazor et al., 2020). Moreover, the interdependencies between perceptual consciousness and monitoring remain to be described empirically: while some theories of consciousness argue that conscious access requires a higher order representation of a stimulus (Brown et al., 2019; Lau & Rosenthal, 2011), or necessarily comes with a sense of confidence (Shea & Frith, 2019), other theories argue that first-order representations may be sufficient (Lamme, 2010; Zeki, 2007). Like for perceptual consciousness, several models propose that evidence accumulation plays an important role for the formation of perceptual confidence (Kvam et al., 2015; Pereira et al., 2020; Pleskac & Busemeyer, 2010; van den Berg et al., 2016). Yet, to our knowledge the underlying neural mechanisms remain to be described.

Here, we sought to investigate the role of evidence accumulation in perceptual consciousness and perceptual monitoring by asking participants to detect weak vibrotactile stimuli and rate their confidence in having detected them. We reasoned that both detection and confidence underlie decision-making processes whereby participants accumulate perceptual evidence over time and gauge its level relative to decision criteria. We examined this possibility in a patient implanted with a microelectrode array in the posterior parietal cortex (PPC, Fig. 1A), considered as one of the functional hotspots of evidence accumulation in the non-human primate brain (Bollimunta & Ditterich, 2012; Gold & Shadlen, 2007; Katz et al., 2016; Roitman & Shadlen, 2002; Zhou & Freedman, 2019). We isolated 368 putative single neurons (Fig. S1) in three different experiments with immediate, delayed, and no-responses in order to characterize the neural correlates of detection and confidence at the single-neuron and population levels and link evidence accumulation in the PPC to perceptual consciousness irrespective of response effectors. These results were generalized in a fourth experiment involving a group of healthy volunteers in whom we recorded scalp electroencephalography, perceptual consciousness and monitoring responses while they detected the same vibrotactile stimuli. In a final step, we test and propose an evidence accumulation computational model that reproduced the behavioral and neural markers of both detection and confidence. Together, these results indicate that subjective reports of perceptual consciousness and monitoring involve a common mechanism of evidence accumulation orchestrated by the PPC.

**Fig.1.**
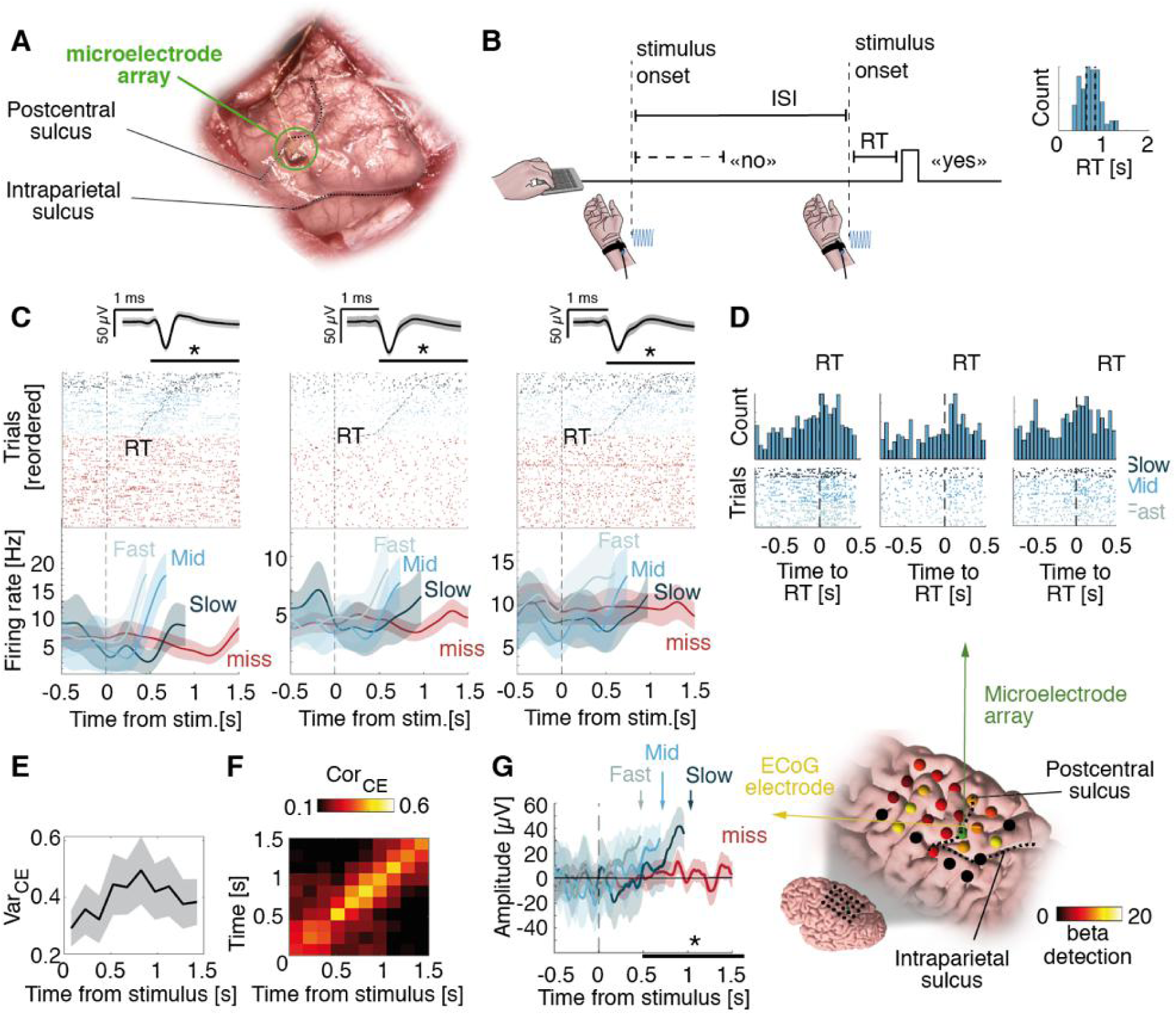
Neuronal correlates of detection in an immediate-response task (Experiment 1). **(A)** Intraoperative photo of the microelectrode array posterior to the postcentral sulcus and dorsal to the intraparietal sulcus. **(B)** The participant pressed a key as soon as he felt a stimulus (dashed vertical lines). In this example, the first stimulus is a miss (i.e. no key press within 2s following stimulus onset) and the second stimulus is a hit. ISI: inter-stimulus interval. Inset: RT distribution. **(C)** Example selective neurons with a latency effect for RT. Top: raster plot time-locked to stimulus onset with spike waveform and shaded standard deviation above. Hits were reordered according to RT (black dashed trace). Bottom: average firing rate for three terciles of RT (blue) and for misses (red). Statistics were performed on continuous data. **(D)** Top: RT-aligned spike count histograms for neurons in **C** (50 ms bins). Bottom: corresponding raster plots. **(E)** VarCE increases during the putative decision process for detection– and RT–selective neurons (N=47). Shaded areas represent 95%-confidence intervals (95%-CI) across selective neurons. **(F)** Corresponding analysis of covariance representing CorCE as heat maps, averaged across detection– and RT–selective neurons (N=47). **(G)** Left: Average ECoG response, aligned to stimulus onset from one electrode posterior to the microelectrode array for three terciles of RT. Right: ECoG grid with beta coefficients for detection. Non-significant electrodes are in black. All shaded areas represent 95%-CI and black horizontal bars represent the analysis window for statistics.

## Results

### Experiment 1: immediate-response task

In *Experiment 1*, the participant was asked to detect vibrotactile stimuli applied to the right wrist (contralateral to the PPC implant) with an intensity around detection threshold. Responses were provided by a keypress with the left hand, immediately after perceiving a stimulus. A trial was considered a hit when the participant responded within 2 s following stimulus onset (41.20% of trials; mean response time (RT) and 95% confidence interval: 0.71 ± 0.02s), otherwise, it was considered a miss (58.80% of trials Fig. 1B). The participant rarely responded “yes” in the absence of stimuli (0.36%; false-alarms), indicative of conservative behavior. We found 94/186 detection-selective neurons (50.54%; p = 0.001, Poisson GLM with permutation test across neurons) with spike counts explained by detection (yes/no responses) between 0.5 to 1.5 s after the stimulus onset. Some neurons were characterized by a hallmark of evidence accumulation where increases in firing rates preceded detection reports depending on their response times (Bollimunta & Ditterich, 2012; Katz et al., 2016; Roitman & Shadlen, 2002; Zhou & Freedman, 2019): the cumulative sum of spikes following stimulus onset correlated with the corresponding RT in 67/94 detection-selective neurons (71.28%; p = 0.001; Fig. 1C) with some neurons showing gradually increasing spike counts prior to the keypress (Fig. 1D). To further support that increased spike counts represent an evidence accumulation process, we verified that the proportion of variance not attributed to the point process increased after stimulus onset (Fig. 1E) and that the corresponding covariance decayed with increasing time lag (Fig. 1F), in line with what is expected for a diffusion process (Churchland et al., 2011). Finally, we replicated the increase in firing rate with electrocorticography (ECoG) by showing that the strongest effect of detection was localized in the PPC and pre-central gyrus (Fig. 1G). To summarize, we uncovered individual neurons in the human PPC with firing rates ramping up prior to detection reports, consistent with evidence accumulation.

### Experiment 2: delayed-response task

Next, we tested whether neuronal responses relate to conscious perception irrespective of motor actions by imposing a delay between stimulus onset and reports. We also assessed whether the strength of these neuronal representations co-vary with reported confidence (Rutishauser et al., 2015, 2018). In Experiment 2, we asked the participant to report vocally the detection of the stimuli with a minimal delay of 1 s after stimulus onset (Fig. 2A, upper panel). To assess the role of evidence accumulation for conscious monitoring, we also asked the participant to vocally report his confidence (high, medium, low) in his response. Similar to Experiment 1, 50.5% of stimuli were detected (hits) (20% trials had no stimuli, of which 5% were false-alarms, confirming his conservative strategy). When a stimulus was presented, confidence was higher following hits (2.46 ± 0.10) than misses (2.00 ± 0.08; X^2^ = 20.09, p = 4.3*10-5), indicative of accurate detection monitoring processes (Fig. 2A, lower panel).

**Fig.2.**
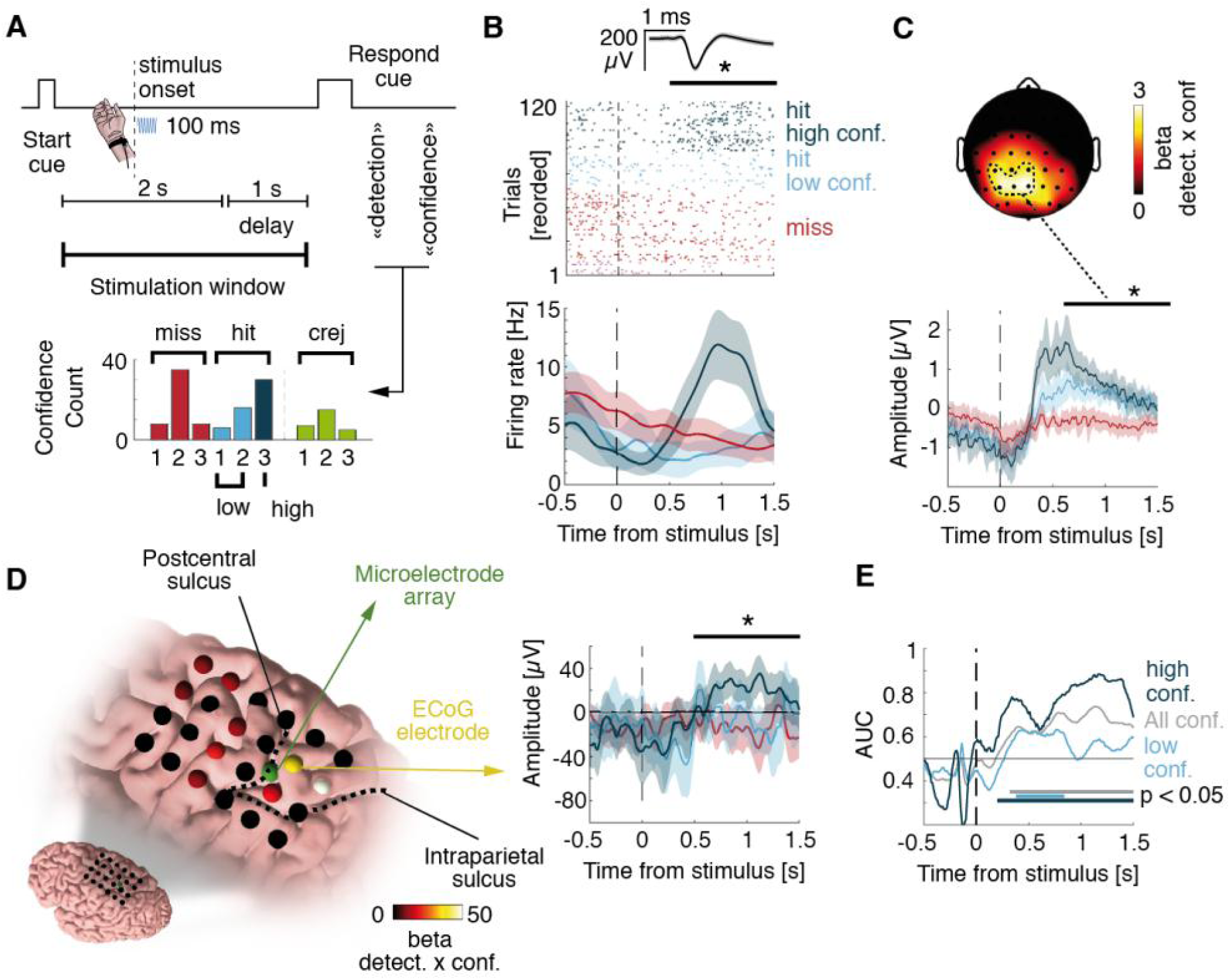
Neuronal correlates of detection and confidence in a delayed-response task (Experiment 2). **(A)** Top: Vibrotactile stimuli were applied during a 2s window following an auditory cue. After 1s delay, the participant was prompted to give detection and confidence reports. Bottom: Distribution of confidence. For display purposes hereafter, signals corresponding to confidence values of 1 and 2 were merged into low-confidence, while confidence values of 3 were considered as high-confidence. Statistics were done on the three levels. **(B)** Example selective neuron. Top: Raster plot time-locked to stimulus onset with spike waveform with shaded standard deviation above. Bottom: Corresponding firing rates. **(C)** EEG data showing a topographic map of beta coefficient for the interaction between detection and confidence for hits (dashed trace). The EEG amplitude time-locked to stimulus onset and averaged over 18 healthy controls is shown below. **(D)** Left: ECoG grid with beta coefficients for detection x confidence. Non-significant electrodes are in black. Right: Average ECoG amplitudes, aligned to stimulus onset from the electrode next to the microelectrode array **(E)** Decoding performance for different confidence levels. Horizontal lines show times of significant performance (permutation tests). All shaded areas represent 95%-CI and black horizontal bars represent the analysis window for statistics.

We ran a factorial analysis to identify neurons encoding detection and/or confidence. We found 17/86 neurons showing an interaction between detection and confidence (19.77%, p = 0.002, permutation test) driven by an increased firing rate for hits with high confidence (Fig. 2B). Only one neuron showed only a main effect of detection (1.16%, p = 0.57) and two a main effect of confidence (2.33%, p = 0.88). A similar interaction between detection and confidence was found in ECoG electrodes surrounding the microelectrode array (Fig. 2D) and in electroencephalography (EEG) signals from 18 healthy volunteers recruited from Experiment 4 (Fig. 2C), consistent with previous EEG studies (Herding et al., 2019; Tagliabue et al., 2019). To characterize how neuronal population activity relates to detection and confidence, we trained decoders on the firing rate of all neurons and evaluated them out-of-sample. We decoded hits from misses better than chance for both high confidence (Fig. 2E; max. area under the curve (AUC): 0.88, 1.16s after stimulus onset) and low confidence (max. AUC: 0.63 accuracy at 0.77 s). This indicates that although low confidence hits and misses were indistinguishable based on individual neurons they could be discriminated at the population-level, which confirms that our results were not driven by high-confidence trials only. Finally, the output of the best decoder (at 1.13 s) correlated with confidence for hits (R = 0.59; p < 0.001, permutation test) but not for misses (R = 0.16; p = 0.13), confirming that the neuronal signal driving detection also explains confidence for detected stimuli. Together, results from Experiments 1 and 2 show that PPC neurons exhibit evidence accumulation behavior and encode detection and confidence reports irrespective of motor actions and report effector (i.e., keypress in Experiment 1, voice in Experiment 2). Of note, the latency of the evidence accumulation process we uncovered in Experiment 2 is qualitatively compatible with the distribution of RTs measured in Experiment 1, which suggests that conscious access occurs with a delay of up to 1 s following weak vibrotactile stimulation.

### Experiment 3: no-report paradigm

To distinguish neuronal activity associated with subjective experience from activity associated with subjective report (Aru et al., 2012; Pitts et al., 2014; Tsuchiya et al., 2015), in *Experiment* 3 we let the participant mind-wander while he was exposed to stimuli ranging between 0.5 to 5 times the perceptual-threshold intensity. We reasoned that neuronal activity would still encode the intensity even for task-irrelevant stimuli if evidence accumulation determines conscious perception beyond mere reports. While no behavioral task was enforced, we found that the activity of 14/96 neurons increased with increasing stimulus intensity (14.58%, p = 0.008; Fig. 3C), similar to hits in Experiments 1-2 (Fig. 3A, B). The fact that stimulus intensity was represented at the single-neuron level although the participant was not engaged in the task argues against the possibility that our previous results in Experiments 1-2 reflected task activity rather than perceptual processing leading to conscious access (Aru et al., 2012; Pitts et al., 2014; Tsuchiya et al., 2015).

**Fig. 3.**
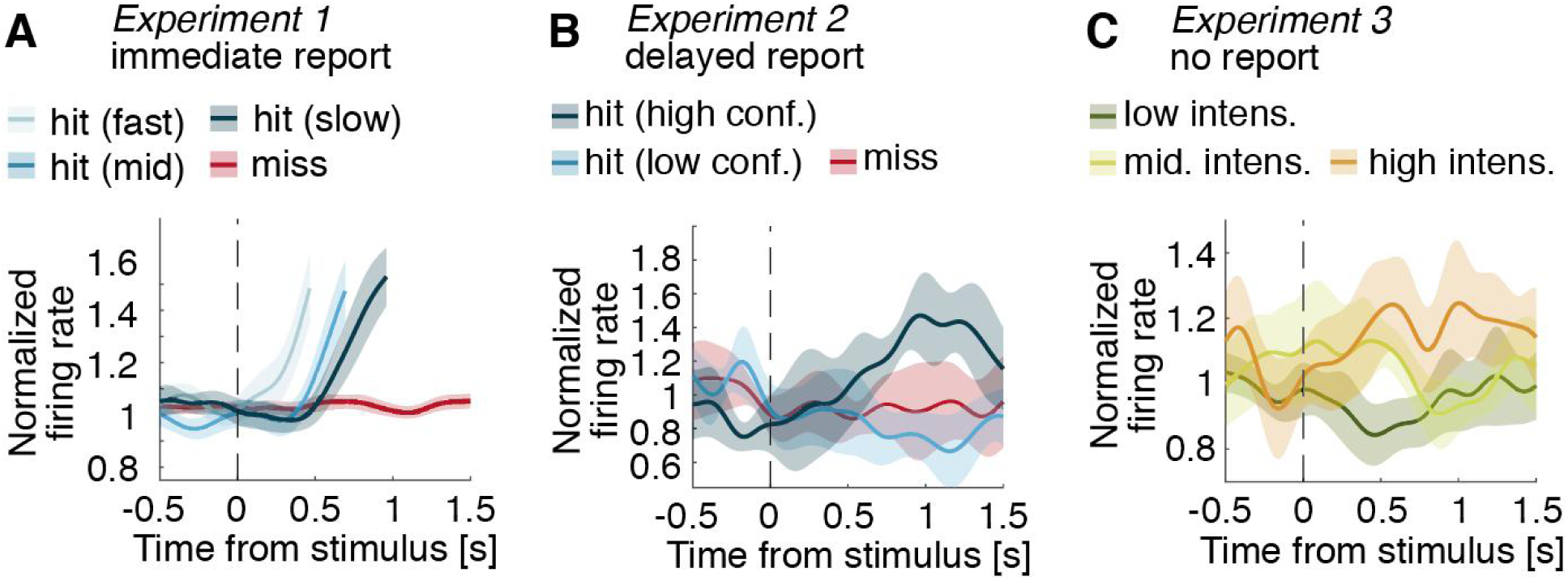
Average firing rates of responsive neurons. Firing rates were normalized using a 0.3 s pre-stimulus baseline. **(A)** Normalized firing rate for three bins of RT (hits; blue) and for misses (red), averaged across detection-selective and RT-selective neurons with higher firing rates for hits (N=47). In Experiment 1, the participant answered with a keypress for hits. **(B)** Normalized firing rate for high and low confidence for hits (blue) and for misses (red), averaged across all detection- and confidence-selective neurons with higher firing rates for hits (N=10). In Experiment 2, the participant waited at least one second before reporting detection and confidence vocally. **(C)** Normalized firing rate for three bins of stimulus intensity, averaged across intensity-selective neurons (N=14). In Experiment 3, the participant provided no detection or confidence report and was let to mind wander.

### Experiment 4: computational model of detection and confidence

Informed by these human single-neuron data from Experiments 1-3, we sought to generalize our decisional account of perceptual consciousness by identifying evidence accumulation mechanisms underlying detection and confidence in EEG data (18 healthy volunteers, task similar to Experiment 2). Participants behaved similarly to the aforementioned patient, with a balanced number of hits and misses (Supplementary results) and EEG responses also showed an interaction effect between detection and confidence (Fig. 2C). We developed an evidence accumulation model to fit the behavioral and EEG data, assuming that participants attempted to detect the stimulus by continuously accumulating evidence during a 3s stimulation window (from trial onset until the response cue). To model the time uncertainty in our task (participants did not know when a stimulus could be applied), we assumed that participants started accumulating evidence before the stimulus onset (Devine et al., 2019). This was modelled as a null drift rate across time except for a short-lasting boost triggered by the stimulus. A stimulus was perceived if the simulated evidence accumulation (EA) process reached a bound (Kang et al., 2017) at any time during the stimulus window (Fig. 4A), compatible with all-or-none views of conscious access (Dehaene et al., 2014).

**Fig. 4.**
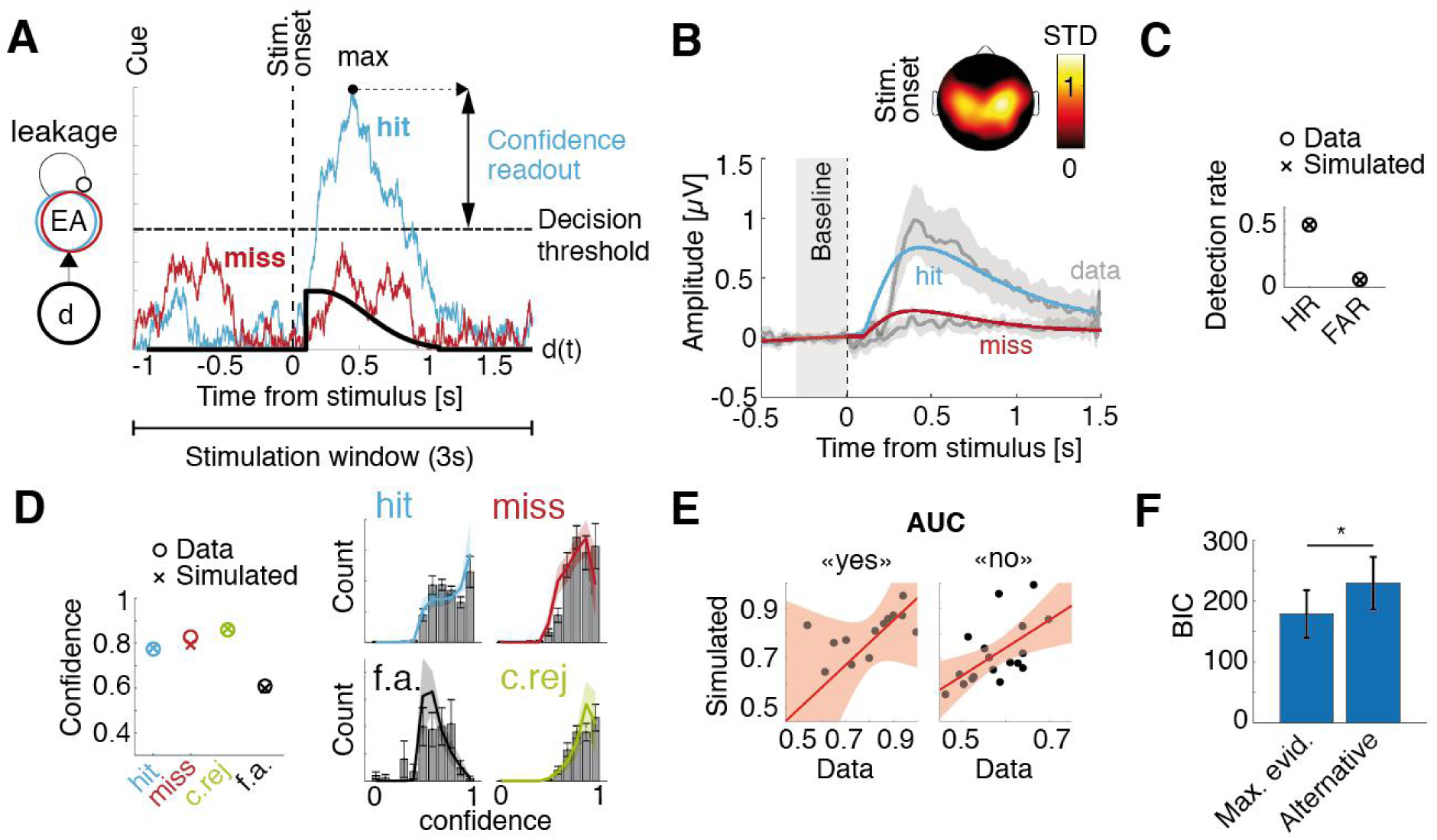
Computational model based on evidence accumulation. **(A)** Time-varying drift rate (d; thick black trace) had a short-lasting boost after a non-decision time following stimulus onset (dashed vertical line). Example evidence accumulation for one trial (EA; cyan trace for a hit, red trace for a miss) rises sharply after the drift boost and is attracted back to zero due to leakage. A stimulus is considered as perceived (hit) if EA reaches a decision threshold (horizontal line), and as non-perceived (miss) if not. The maximum of accumulated evidence with respect to the decision threshold is used as a confidence readout. **(B)** Model fit of the pEA locked on stimulus onset for hits (cyan trace) and misses (red trace). The corresponding observed EEG data is shown in grey. Average scalp topography of pEA weights is shown above. **(C)** Hit rate (HR) and false alarm rate (FAR). Datapoints are represented as ‘o’ and model simulations as ‘x’. **(D)** Left: Average confidence for hits (cyan), misses (red), correct rejections (green) and false alarms (black). Right: Model fits of the confidence distributions. Histograms show confidence distributions with 95%-CI whiskers. Colored traces show model simulations. All shaded areas represent 95%-CI. **(E)**. Area under the curve (AUC) correlation between observed data (horizontal axis) and simulated data (vertical axis) for “yes” responses (hits and false alarms; left) and “no” responses (correct rejections and misses; right). Regression line is shown in red with shaded areas representing 95%-CI). (**F)**. Model comparison in terms of Bayesian information criterion (BIC) between the maximal evidence model and the alternative model. Whiskers represent 95%-CI and asterisk indicates statistical significance (p<0.05).

Confidence was read out from the distance between accumulated evidence and the decision threshold (Pereira et al., 2020; Pleskac & Busemeyer, 2010). Importantly, we sampled confidence when evidence reached a maximum across the stimulation window, which allowed implementing a confidence readout for misses and correct rejections, for which no decision threshold is crossed. To fit the model parameters to the data, we considered the shape of the electrophysiological signature for hits and misses as a neural correlate of evidence accumulation (O’Connell et al., 2012; Philiastides et al., 2014; Tagliabue et al., 2019), defined by the weighted average of all EEG electrodes that maximally discriminated hits from misses. We first fitted the parameters of a detection model to these electrophysiological responses (Fig. 4B, S2) as well as to hit and false alarm rates (Fig. 4B, inset; Fig. S3). We then fitted two additional parameters for confidence bias and sensitivity to observed confidence distributions. The resulting model fitted the confidence ratings well (average R across participants 0.83±0.03 for hits, 0.85±0.03 for misses, 0.81±0.04 for correct rejections and 0.45±0.09 for false alarms; Fig. 4C, S4), suggesting that evidence accumulation is a plausible mechanism underlying perceptual consciousness and its electrophysiological correlates. The data and the model were still consistent when stratifying per confidence level. Metacognitive sensitivity predicted by our model and observed in the data were correlated for both “yes” responses (R=0.60, p=0.001, permutation test) and “no” responses (R=0.61, p=0.009; Fig. 4E), showing that our model also successfully predicted metacognitive performance. Finally, an alternative model assuming that confidence for detected stimuli is sampled at a fixed latency after crossing the decision threshold and confidence for undetected stimuli is sampled from a random distribution led to a worse fit of the data (BIC = 229.31 ± 43.19 compared to BIC = 178.82 ± 38.86 for the maximal evidence model; z = -2.33, p = 0.020; Fig. 4F). This difference in goodness of fit was also observable in the correspondence between observed and simulated averaged confidence for hits (R = 0.81 ± 0.03 for the maximal evidence model compared to R = 0.63 ± 0.09 for the alternative model; z = 2.29; p = 0.022).

Informed by our modelling results in healthy participants, we set out to verify whether we could decode confidence for misses from single-neuron data in Experiment 2 using a decoder defining confidence as the maximum of accumulated evidence. We took the best decoder for hits and misses (trained on stimulus-locked data) and applied it out-of-sample across the stimulation window (i.e. cue-locked). We decoded confidence for hits (R = 0.49, p = 0.001, permutation test) and confidence for misses (R = -0.31, p = 0.015), which the aforementioned stimulus-locked decoder could not achieve. The time corresponding to the decoded maximal evidence correlated with stimulus onset for hits (R = 0.37, p = 0.001) but not for misses (R = 0.03, p = 0.58), suggesting that evidence for confidence in misses was not sampled synchronously with the stimulus, thereby verifying the plausibility of the maximal evidence decoder on our patient’s single-neuron data.

## Discussion

We propose a mechanism of evidence accumulation to explain the behavioral and neural markers of perceptual consciousness and monitoring. We show that tactile detection relates to an increase of the firing rate of single neurons in the posterior parietal cortex of a human participant, as well as an increased scalp EEG response recorded in a group of healthy participants. In both cases, the amplitude of the corresponding neural response was dependent on the confidence in hits. This increase in neural response as well as in the detection reports were well described by a computational model indicating that a plausible mechanism underlying the building of confidence in both the presence and absence of a stimulus is for the brain to take the maximal evidence accumulated over time.

### Encoding of detection by individual neurons in the posterior parietal cortex

We had the opportunity to collect data from individual neurons in the human PPC, at the junction between the postcentral and intraparietal sulcus in the superior parietal lobule. The PPC has been associated with a multitude of functions linking perception to planning and action (Andersen & Cui, 2009) and receives multisensory inputs including those from the primary somatosensory cortex (Pearson & Powell, 1985). In Experiment 1, we found individual neurons in the PPC with higher firing rates following detected stimuli. We argue that these neurons are responsible for evidence accumulation based on the following three findings. Firstly, in Experiment 1 we found neurons whose increase in firing rate for hits was synchronized to response times (Gold & Shadlen, 2007). Secondly, in Experiment 1 the variance of the corresponding spike rates increased after stimulus onset (Churchland et al., 2011). Thirdly, in Experiment 3, we observed an increase in the firing rates for increasing intensities of (task-irrelevant) stimuli (Gold & Shadlen, 2007). Based on these three hallmarks of evidence accumulation, we argue that our results consist in the first single-neuron account of perceptual evidence accumulation in a human subject capable of subjective reports. Indeed, although electrophysiological correlates of evidence accumulation have been found in various regions of non-human primate brains, including the frontal cortex or subcortical structures (Ding & Gold, 2010; Hanks et al., 2015; Odegaard et al., 2018), the arguably most common region studied in relation with neural accumulation of perceptual evidence is the lateral intraparietal (LIP) area of the PPC (Bollimunta & Ditterich, 2012; Katz et al., 2016; Roitman & Shadlen, 2002; Zhou & Freedman, 2019). However, whether perceptual evidence accumulation neurons such as those reported in non-human primate studies could support conscious reports is unclear, as subjective experience cannot be measured explicitly in non-human species, and because such neurons were – to our knowledge – not reported in humans yet.

### Encoding of confidence by individual neurons in the posterior parietal cortex

In Experiment 2, we asked the participant again to detect stimuli and found neurons similar to those in Experiment 1 with higher firing rates after stimuli reported as perceived. The finding of detection-selective neurons when responses were provided by key press (Experiment 1) or orally (Experiment 2) suggests that the mechanism of evidence accumulation we propose is response-invariant. Importantly, we asked the participant to report the confidence he had in his responses, and found that the change in firing rates for detected stimuli was modulated by confidence, showing that confidence relates to the strength of single neuron’s responses to detected stimuli. This mechanistic overlap, which – to our knowledge – was not yet shown in humans capable of reporting subjective confidence was confirmed at the neuronal population level: multivariate decoders trained to discriminate hits vs. misses allowed us to decode confidence for hits when time-locking to the stimulus onset and for both hits and misses when locking on the onset of maximal evidence.

### Computational modelling and replication at the scalp level

Because microelectrode implants in parietal regions are extremely rare in humans, we sought to generalize our findings by recording behavioral and neural data in a group of healthy volunteers in Experiment 4. Behavioral results revealed highly similar patterns between the two samples, indicating that detection and confidence reports in Experiments 1-2 were not impacted by the clinical condition. Experiment 4 also allowed us to generalize our single-neuron findings at a larger scale based on EEG recordings. EEG data showed a similar dependence of detection-related activity on confidence for hits, similar to previous work in the visual domain using a different awareness scale (Tagliabue et al., 2019) and a discrimination task (Boldt & Yeung, 2015; Gherman & Philiastides, 2015). Of note, both neural responses recorded at the intracranial (single-neuron, EcoG; Experiment 1-3) and scalp levels (Experiment 4) were observed at a rather late latency following stimulus onset (>200 ms), suggesting that these responses were not related to early somatosensory perceptual processes. We then reasoned that since similar increases in neural activity are assumed to reflect accumulation of evidence (O’Connell et al., 2012, 2018; Tagliabue et al., 2019), an evidence accumulation model should predict both behavioral results (hit rate, false-alarm rate and confidence for hits, misses, correct rejections and false-alarms) and corresponding neural responses. Using neural data to fit the model instead of response times allowed us to fit a leakage parameter (Yu et al., 2015) and to compensate for the fact that, in a detection task, response times are unavailable for undetected stimuli. The model derives confidence as the distance between the maximal evidence accumulated over time and the decision bound. This flexible readout provides a major advantage when computing confidence in the absence of stimulus detection (i.e. misses) as well as in the absence of a stimulus (i.e. correct rejections and false-alarms), which cannot be achieved with models using decision-locked confidence readouts in discrimination tasks (Pereira et al., 2020; Pleskac & Busemeyer, 2010; van den Berg et al., 2016). An alternative model with a fixed-timing readout and a random confidence for unperceived stimuli performed significantly worse. Our modelling results corroborate our electrophysiological results across Experiments 1-4 and are consistent with decision-making models of confidence applied to animal data, postulating a shared encoding of evidence for decision and confidence (Kiani & Shadlen, 2009), possibly enriched by post-decisional evidence (Fleming et al., 2018; Pleskac & Busemeyer, 2010; van den Berg et al., 2016). Moreover, a recent study comparing models based on signal detection theory showed that the model that best fit observed data involves a second-order “metacognitive” noise to the decisional evidence (Maniscalco & Lau, 2016). Our model implicitly implements this metacognitive noise through the influence of first-order parameters such as leakage on post-decisional evidence readouts. Indeed, in participant with strong leakage, accumulated evidence rises and decays fast, leading to low metacognitive noise. On the contrary, in participants with little leakage, once no more informative evidence is accumulated, the level of evidence accumulation tends to oscillate around the reached maximum, leading to higher metacognitive noise. A-posteriori analyses showed that the leakage parameter correlated with metacognitive sensitivity (Supplementary information). Our model thus provides a simple mechanism supporting metacognition, including subliminal stimuli, explains neural responses and was verified at the single neuron and EEG level, as we were able to decode confidence ratings above chance using this procedure.

### Implications for perceptual consciousness

Our results posit that stimulus detectability involves the accumulation of sampled evidence towards a decision bound, as previously discussed for discrimination tasks (Kang et al., 2017). Of note, the use of a detection task is compatible with a contrastive study of consciousness (Baars, 1998), as opposed to two-alternative forced choice discrimination tasks for which confidence ratings are well characterized, but which do not offer a direct contrast between perceived and unperceived stimuli. The view of conscious access as an all-or-none process involving a decision bound is compatible with the ignition mechanism put forward by the global workspace theory of consciousness (Mashour et al., 2020), according to which sufficiently activated encapsulated networks may coalesce into a single network responsible for broadcasting neural signals throughout the brain and thereby making them accessible to conscious reports. One could speculate that the triggering of an ignition is governed by bounded-evidence accumulation similar to the one operated by neuronal populations in the posterior parietal cortex. Recently, the use of classical contrastive approaches to delineate the neural correlates of consciousness has been criticized, on the basis that it may be confounding the cognitive and neural mechanisms associated with phenomenal experience per se, and those associated with reporting phenomenal experience (Aru et al., 2012). Some authors have proposed the use of “no-report paradigms”, in which perceptual experience is not inferred from participants’ responses, but from neural or peripheral signals while participants are passively exposed to stimuli (Frassle et al., 2014; Tsuchiya et al., 2015; but see Block, 2019; Phillips & Morales, 2020). Importantly, we found a population of neurons encoding perceptual evidence through a putative evidence accumulation process in Experiment 3, in which the participant was passively exposed to the stimuli similar to such no-report paradigms. Although these effects were weaker than the ones found in Experiment 1-2, they indicate that evidence accumulation operated by a neuronal population in the posterior parietal cortex is involved in conscious perception, even when the stimuli are task-irrelevant. In addition, the mechanistic overlap between detection and confidence we report cannot be due to similar motor responses associated with detection and confidence reports in Experiment 2, as those were collected separately, seconds after the end of the stimulation window on which our analysis was based in the delayed detection task.

To conclude, our results posit that both detection and confidence for near-threshold stimuli involve the accumulation of evidence towards a criterion orchestrated by the PPC. We argue that this neuronal mechanism involving a decision bound may serve as a trigger for the neural ignition underlying conscious access (Moutard et al., 2015; van Vugt et al., 2018) and explains how contents remaining inaccessible to consciousness may still be subject to self-monitoring (Mazor et al., 2020; Meuwese et al., 2014). Our behavioral, neural, and modeling results clarify how perceptual consciousness and reflexive self-monitoring are intertwined mechanistically.

## Supporting information

Supplementary results

## STAR METHODS

### KEY RESSOURCE TABLE

| REAGENT or RESSOURCE | SOURCE | IDENTIFIER |
| --- | --- | --- |
| **Software and Algorithms** |  |  |
| **MATLAB 2018a** | MathWorks | RRID:SCR_001622;<br>http://www.mathworks.com/products/matlab/ |
| **Osort** |  | RRID:SCR_015869;<br>http://www.rutishauserlab.org/osort |
| **EEGLAB 2019.1.0** |  | RRID:SCR_007292;<br>http://sccn.ucsd.edu/eeglab/index.html |
| **iELVis** |  | RRID:SCR_016109;<br>http://ielvis.pbworks.com/w/page/116347253/FrontPage |
| **Psychophysics toolbox 3** |  | RRID:SCR_002881; http://psychtoolbox.org/ |
| **Algorithm to fit an evidence accumulation model to behavioral and neural data** |  | Upon acceptance |
| **Other** |  |  |
| **Neuroport recording system** | Blackrock Microsystems | https://www.blackrockmicro.com/ |
| **Brain Quick LTM** | Micromed | http://www.micromed.eu/en-us/ |
| **WaveGuard EEG** | ANTNeuro | http://www.ant-neuro.com |

### CONTACT FOR REAGENT AND RESSOURCE SHARING

Further information and requests for ressources should be addressed directly to the Lead Contact, Nathan Faivre

### EXPERIMENTAL MODEL AND SUBJECT DETAILS

In experiments 1–3, the participant was a 23-year-old right-handed man suffering from drug-resistant epilepsy due to a focal cortical dysplasia in the left central sulcus. As part of the clinical management of his condition, he received a 4×6 ECoG grid covering the left premotor, motor, sensory and posterior parietal cortices. He accepted to participate in a clinical trial on neuronal recordings during invasive epilepsy monitoring at the Geneva University Hospitals (IN-MAP; NCT02932839) and a Utah microelectrode array was additionally implanted in the left posterior parietal cortex. The patient provided informed written consent and the study was approved by the Commission Cantonale d’Ethique de la Recherche de la République et Canton de Genève (2016-01856). Eighteen healthy participants (7 females; age: 25.2 years, SD = 4.1) took part in Experiment 4 for a monetary compensation. Participants gave written informed consent prior to participating and all experimental procedures were approved by the Commission Cantonale d’Ethique de la Recherche de la République et Canton de Genève (2015-00092 15-273).

### METHOD DETAILS

Experimental paradigms were written in Matlab (Mathworks) using the Psychophysics toolbox (Brainard, 1997; Kleiner, n.d.; Pelli, 1997). In all experiments, stimuli were applied on the lateral palmar side of the right wrist using a MMC3 Haptuator vibrotactile device from TactileLabs Inc. (Montréal, Canada) driven by a 230 Hz sinusoid audio signal lasting 100 ms. Experiments started by a simple estimation of the individual detection threshold. The tactile stimulus was applied with decreasing intensity with steps corresponding to 2% of the initial intensity until the participant reported not feeling it anymore three times in a row. We then repeated the same procedure but with increasing intensity and until the participant reported feeling the vibration three times in a row. The perceptual threshold was estimated to be the average between the two thresholds found using this procedure. This approximation was then used as a seed value for an adaptive staircase during the main experiments (see below). Experiments 1-3 were performed on different days at the patient’s bedside.

#### Experiment 1

Stimuli were applied in a pseudo-random way with an inter-stimulus interval of two seconds plus an exponentially distributed time (mean: 2 s). The participant was provided with a keypad and asked to press a key every time he felt a stimulus. Answers provided during the two seconds following a stimulus were considered as hits. Only one keypress occurred out of this two second post-stimulus window.

#### Experiment 2

An auditory cue signaled the start of the two seconds stimulus window during which the stimulus could be applied at any time (uniform distribution) in 80% of trials (the remaining 20% served as catch trials, unbeknownst to the participant). Stimulus onset was followed by a one second delay to ensure that stimulus-locked activity was not contaminated by the detection response. After this delay, a second auditory cue probed the participant for his detection response (“yes” or “no”), followed by a three levels confidence rating (1: “unsure”, 2: “somewhat sure”, 3: “very sure”). Detection and confidence ratings were provided vocally and registered by the experimenter.

#### Experiment 3

Stimuli were applied in a pseudo-random way with an inter-stimulus interval of two seconds plus an exponentially distributed time (mean: 2 s) with a random amplitude sampled from 11 intensities ranging from zero to five times the participant’s perceptual threshold. The participant was not given any instructions and was left free to mind-wander during the experiment.

#### Experiment 4

Participants sat in front of a computer screen. A white fixation cross appeared in the middle of the screen for 2 s. From the moment the fixation cross turned green, participants were told that a tactile stimulus could be applied at any moment during the next 2 s. During this period, stimulus onset was uniformly distributed in 80% of trials, the 20% remaining trials served as catch trials, as in Experiment 2. In all trials, 1 second after the green cross disappeared, participants were prompted to answer with the keyboard whether they felt the stimulus or not. Following a 500 ms stimulus onset asynchrony, participants were asked to report the confidence in their first order response by moving a slider on a visual analog scale with marks at 0 (certainty that the first-order response was erroneous), 0.5 (unsure about the first-order response) and 1.0 (certainty that the first-order response was correct). Detection and confidence reports were provided with the left (non-stimulated) hand, using different keys. The total experiment included 500 trials divided in 10 blocks, and lasted about 2 hours.

#### Electrophysiological data acquisition

A 96-channel silicon-based microelectrode array (“Utah array”; Blackrock Microsystems, Salt Lake City, USA) was implanted in the posterior parietal cortex, immediately posterior to the postcentral sulcus and the hand representation of sensorimotor cortex (Fig. 1A). The location was confirmed through post-hoc electrode localization (Fig. 1G), performed through a coregistration of a preoperative MRI structural T1 scan and a postoperative CT scan using the iELVIS toolbox (Groppe et al., 2017). The data from each of the 96 channels was amplified and sampled at 30 KHz for offline analysis (NeuroPort system, Blackrock Microsystems LLC, Salt Lake City, USA). Additionally, a 24 electrode ECoG grid (Ad-Tech Medical) covered the left hemisphere from the premotor cortex to the superior parietal lobule (Fig.1G, 2D). The data was amplified and sampled at 2048 Hz (Brain Quick LTM, Micromed, Treviso, Italy). In Experiment 4, electroencephalographic data were acquired from 62 active electrodes (10-20 montage) using a WaveGuard EEG cap and amplifier (ANTNeuro, Hengelo, The Netherlands) and digitized at a sampling rate of 1024 Hz. Horizontal and vertical electrooculography (EOG) was derived using bipolar referenced electrodes placed around participants’ eyes. The audio signal driving the vibrotactile actuator was recorded as an extra channel to precisely realign data to stimulus onset.

### QUANTIFICATION AND STATISTICAL ANALYSIS

#### Invasive electrophysiological data processing and univariate analysis

The raw signal from the microelectrode array was bandpass filtered between 300 and 3000 Hz for spike sorting. Trials with epileptic activity or other artifacts were removed from further analysis following visual inspection of ECoG data. Spikes were extracted and sorted using the semi-automatic template matching ‘Osort’ algorithm (Rutishauser et al., 2006). Standard quality metrics were computed for each putative single unit in order to assess their quality (Fig. S1). We computed the firing rate every 1ms with a 100 ms standard deviation Gaussian sliding window.

In Experiment 1, a neuron was considered detection-selective when a significant (two-sided test) effect of detection was found on the number of spikes during a time window between 0.5 and 1.5 seconds after stimulus onset using a generalized linear model (GLM) with a Poisson distribution (Fu et al., 2019; Rutishauser et al., 2018). For this, we fitted a model with one beta regressor: #spikes ∼ β0 + β1*detection (hit or miss). For the latency analysis, we computed the cumulative sum of spikes starting at stimulus onset and correlated (Spearman) it with RTs for every time step (1 ms) between 0 and 1.5 s after stimulus onset. A neuron was considered RT-selective if the correlation was significant within this time range after correcting for false-discovery rate (FDR). To ensure that there was no overfitting and that our results were not driven by outliers, we used a non-parametric permutation test to assess whether the number of selective neurons was significantly above chance; we repeatedly (N=1000) applied the same tests on shuffled data and counted the number of selective neurons. We defined the p-value as the proportion of times that the number of selective neurons for shuffled data was higher than the number of selective neurons found in the data (Fu et al., 2019; Kaminski et al., 2017; Rutishauser et al., 2018). When no selective neuron was found in the shuffled data, we set p = 1/N = 0.001. In Experiment 2, we also used a Poisson GLM to regress the number of spikes during a time window between 0.5 and 1.5 seconds after stimulus onset. We fitted a model with three beta regressors: #spikes ∼ β0 + β1*detection + β2*confidence + β3*detection*confidence. We only interpreted main effects (β1, β2) in the absence of interactions (β3). If β3 was significant (two-sided test), we considered the neuron as detection- and confidence-selective. If β1 was significant but β3 was not, we considered the neuron only detection-selective (idem for confidence-selective). We applied the same permutation test as for Experiment 1. In Experiment 3, we fitted a Poisson GLM with one beta regressors: #spikes ∼ β0 + β1*intensity to find intensity-selective neurons showing an increased firing rate with increasing stimulus intensity and applied the same permutation test as for Experiment 1 and 2.

For ECoG analyses, we re-referenced the channels to a common average and applied a lowpass filter (Hamming window with a cutoff frequency of 40 Hz). Trials with epileptic activity or other artifacts were removed from further analysis following visual inspection. We used linear models (LM) for statistics using the same regressors as for spike counts (see above). For display purposes only, we additionally smoothed the data with a 200 ms Savitzky-Golay filter (Savitzky & Golay, 1964).

#### Analysis of variance for Experiment 1

To further relate our results to evidence accumulation, we analyzed some typical signatures of drift diffusion-like processes: the variance in the number of spikes during an epoch can be decomposed into i) the variance of the point process expected if the neuron had a constant firing rate across trials, and ii) the remaining variance of the conditional expectation (VarCE) which is due to the variability of the neuron’s underlying firing rate across trials. If neuronal activity follows a diffusion process, then VarCE should increase with decision time. Similarly, the correlation between conditional expectations (CorCE) should be stronger between adjacent time windows and decrease with time lag between time windows (Churchland et al., 2011). We followed the approach in Churchland and colleagues. In brief, we relied on an upper bound estimate of VarCE, assuming that the point process variance is proportional to the spike count so:

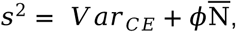

with *s*^*2*^ the total variance and 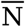 the average number of spikes for one 50 ms epoch. We used 10 non-overlapping epochs ranging from 0 to 1.5 s post-stimulus. The variance of the point process was computed as is a weighted average of the variance of the point process for hits and for misses. The constant *ϕ*can be set based on some heuristics such as by considering that VarCE is zero when the ratio of variance and mean firing rate is minimal (e.g. at the beginning of the decision process). Due to the limited number of trials available in this study, we preferred this approach to more complex methods involving data fitting.

We computed CorCE by dividing the covariance of the number of spikes for different epochs by the square root of the product of the VarCE:

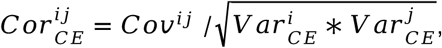

with *Cov*^*ij*^ the covariance between epoch i and j and 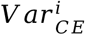 the VarCE for epoch i. For all computations, we reduced *<* by 20% to have positive semi-definite covariance matrices.

#### Multivariate decoding (Experiment 2)

We fed single neurons firing rates sampled every 10 ms into linear discriminant decoders with a L2 regularization factor of 0.8 (similar results were obtained with different regularization factors). These decoders separated the space of input features with a linear hyperplane that best discriminates hits and misses. The decoders predict a hit when the distance of a sample to the separating hyperplane is higher than zero and a miss otherwise. To avoid overfitting, we separated our data in 10 cross-validation folds so that for each fold we trained decoders on the 90% of the data and tested them on the 10% remaining (i.e. out-of-sample). The distance to the separating hyperplane was also used to compute the area under the curve (AUC) for out-of-sample data at each time point. We used permutation tests to assess whether similar AUC could have been obtained by chance while correcting for multiple comparisons across time points. We selected contiguous clusters of time points when AUC was higher than 0.6 or lower than 0.4 and then computed the proportion of similar or bigger clusters obtained with shuffled labels. For each permutation, we computed AUC clusters over the whole set of time points to keep the autocorrelation structure in the shuffled data.

Confidence was decoded at the time point corresponding to the highest AUC. Decoded confidence was defined as the absolute distance of a sample to the separating hyperplane. We assessed confidence decoding by correlating (Spearman) decoded confidence with observed confidence and assessing significance using permutation tests (one-sided, as we could reasonably expect positive correlations). As this procedure was carried out on out-of-sample data, our results were not affected by overfitting. Finally, for maximal evidence decoding, we used the same decoder to decode firing rates between 0.5 and 3 s post-cue. We excluded the first 0.5 s due to some neurons showing post-cue activity. We then took the maximum of the decoder over that time window, correlated it with confidence observed in the data and assessed significance using permutation tests (one-sided). Again, this procedure was carried out on out-of-sample data so our results cannot be affected by overfitting.

#### Scalp EEG data preprocessing (Experiment 4)

All channels were high-pass filtered using a Hamming window with a cutoff frequency of 0.1 Hz. We defined an epoch as the 3 seconds of data centered around the event corresponding to the vibrotactile stimuli recorded using an auxiliary channel. EEG and EOG data were then lowpass filtered using a Hamming window with a cutoff frequency of 40 Hz and visually inspected to remove trials and channels containing artifacts. We computed the independent component analysis (ICA) (Makeig et al., 1996) on a copy of the EEG epochs that were highpass filtered at 1Hz. The number of independent components (ICs) computed corresponded to 99% of the variance, which resulted in 14.78 ± 1.27 ICs per subject. We used SASICA (Chaumon et al., 2015) with default parameters to automatically select ICs for rejection and visually inspected all components scheduled for rejection before actually rejecting them. IC weights kept were then back-projected to the original EEG epochs. Any channel rejected prior to the ICA was reinterpolated using spherical interpolation (N = 0.67 ± 0.23, max 3). Finally, we visually inspected all channels and rejected artifactual epochs. All pre-processing was done with the EEGLAB toolbox (Delorme & Makeig, 2004). The final dataset comprised 464.17 ± 13.73 epochs per subject.

To assess which electrodes were detection- and/or confidence-selective, we used a linear mixed model to regress single-trial average EEG responses in the 0.5 to 1.5 time-window used for single-neuron analysis. For each electrode, we tried different random factors and kept the model with the lowest Bayesian Information Criterion. P-values were Bonferroni corrected for multiple comparisons. To compute the electrophysiological correlate of evidence accumulation, we sought to find the best weighting of EEG electrodes in terms of discriminability between hits and misses. For this, we trained decoders of hits versus misses between -0.5 and 1.5 s from stimulus onset with 20 ms steps in a 10-fold cross-validation scheme. We repeated this procedure 10 times and searched for the time point with highest average discriminability. We then retrained one decoder using the EEG at this time point and used the weights of this decoder to construct one single value at every time point, representing a proxy to the amount of evidence for hits. We baselined this proxy signal using the 300 ms pre-stimulus.

#### Ornstein-Uhlenbeck bounded accumulation model

To test whether the observed electrophysiological correlate of conscious detection could index evidence accumulation, we used an Ornstein–Uhlenbeck process which consists of a drift diffusion model with a leakage parameter driving the accumulated evidence back to zero (Busemeyer & Townsend, 1993). The model consisted of an evidence accumulation process EA(t) integrating a time-varying drift rate d(t) with a leakage factor *λ* plus additive white noise W(t) with a fixed standard deviation of *σ* = 0.1 (Eq. 1). The evidence accumulation process EA(t) was bounded by zero to be more biologically plausible (since firing rates are positive).

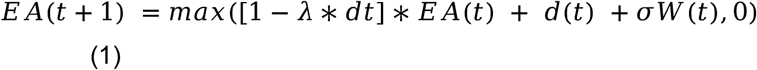

To model temporal uncertainty, the accumulation process started at the beginning of the stimulus window (Devine et al., 2019) with a drift of zero and ended 3 s later, as in our experimental paradigm. On stimulus onset, and after a non-decision time (*ndt*), the drift rate d(t) rose to a level *γ* for 100 ms (stimulus duration) and then decayed exponentially with a factor *k* (Eq. 2). We used the same distribution of stimulus onsets as in the data (from 0 to 2 s after the cue). To model variability in the drift rate for a detection task (Ratcliff & Van Dongen, 2011), *γ* was sampled from a half-normal distribution with mean *γ* _ μand standard deviation *γ*_ *σ* (i.e. absolute value of a normal distribution). This allowed us to have variability in the drift rate while keeping it positive.

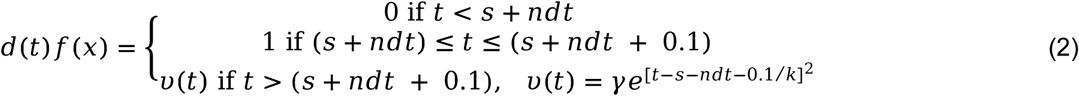

With s the onset of the stimulus. Stimuli were considered to be detected if the accumulated evidence reached a decision bound *θ*.

Confidence c(t) was simulated as the accumulated evidence EA(t), scaled by a factor *α* and shifted by a factor *β*(confidence bias), inverted if the decision bound was not reached (invert(x) = 1 - x) before being saturated to the 0 - 1 interval by a sigmoidal function (Pereira et al., 2020) (Eq. 3). The time at which confidence c(t) was read-out corresponded to the maximum of EA(t) over the 3s stimulation time window. Of note, we expressed the confidence readout in percentage of the decision bound. This scaling did not affect simulated confidence but normalized the readout across participants and helped restrain the grid search for good initial parameters.

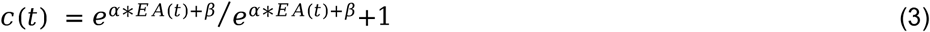

#### Model fitting

We used a two-stage fitting procedure: We first fitted the parameters of the decisional process to detection responses and to the pEA described above and then fitted a second set of parameters to predict confidence ratings.

In the first stage, we simulated (N=500) trials of EA(t) along with the corresponding detection responses. The objective function of the optimization procedure was based on the likelihood of the model with respect to the hit rate and false-alarm rate observed in the data and the shape of the average of pEA(t) for hits and misses. Since our model is agnostic to the scale of pEA(t) and EA(t), we scaled them both by their average over time and realizations (trials or simulations). The log-likelihood thus corresponded to the normal probability of observing such a hit-rate and false-alarm rate and the normal probability of observing such an electrophysiological response for hits and misses. We used a Nelder-Mead simplex optimization with 756 different initial parameters sampling a broad range of values for *γ_ μ, σ*_ *γ, λ* and *k*. For each such iteration, we first did a grid search on ndt and *θ* to find plausible starting values. We kept the parameters corresponding to the model with the best likelihood. To ensure a good fit, we did a final fitting with N=10’000 simulated trials.

In the second stage, we also simulated (N=500) trials of EA(t) along with the corresponding detection responses and used these to simulate confidence ratings. We used a Kolmogorov-Smirnov test for the log-likelihood of confidence for hits, misses and correct rejections. We used a Nelder-Mead simplex optimization with 66 different initial parameters sampling a broad range of values for *α* and *β*. We kept the parameters corresponding to the model with the best likelihood. To ensure a good fit, we did a final fitting with N=10’000 simulated trials.

#### Metacognitive sensitivity

We evaluated metacognitive sensitivity or how well confidence predicted task performance (Fleming & Lau, 2014). For this, we assessed the relation between confidence ratings and detection accuracy using the area under the curve, computed independently for “yes” responses (hits and false alarms) and “no” responses (correct rejections and misses) (Mazor et al., 2020; Meuwese et al., 2014)

#### Alternative model

We compared our maximal evidence model with an alternative model which assumed that confidence was readout at a fixed timing (t_RO) post-decision for perceived stimuli. For unperceived stimuli (i.e. with no decision), this alternative model assumed that participants reported random confidence estimates, based on Gaussian noise with mean Ω and unit standard deviation. As in the maximal evidence model, confidence corresponded to evidence scaled by a factor α and shifted by a factor β (confidence bias) before being saturated to the 0 - 1 interval by a sigmoidal function. The model thus comprised four parameters: α, β, t_RO and Ω which were fitted with the same procedure as for the maximal evidence model, except that the grid search for optimal initial parameters was extended to include two initial values for t_RO: 0.25 and 0.5 s. We used the Bayesian Information Criterion to compare the two models.

## DATA AND SOFTWARE AVAILABILITY

Behavioral, electrocorticographic and electroencephalographic data with the corresponding analyses scripts will be made available upon publication. Single-neuron data are available upon request.

## Acknowledgments

MP is supported by an Early Postdoc.Mobility fellowship from the Swiss National Science Foundation (P2ELP3_187974). PM is supported by an Ambizione fellowship from the Swiss National Science Foundation (PZ00P3_167836). MS is supported by grants from the Swiss National Science Foundation (163398, 80365) and Fondation Privé. OB is supported by the Bertarelli Foundation, the Swiss National Science Foundation, and the European Science Foundation. NF has received funding from the European Research Council (ERC) under the European Union’s Horizon 2020 research and innovation programme (Grant agreement No. 803122). We acknowledge support from the Wyss Center for Bio and Neuroengineering and the Epileptology Unit, Division of Neurology, Geneva University Hospitals. We thank Jean-Baptiste Eichenlaub, Vincent de Gardelle, Emanuela de Falco, Liad Mudrik and Adam Shai for their comments.

## The authors declare no competing interests

## References

Andersen, R. A., & Cui, H. (2009). Intention, Action Planning, and Decision Making in Parietal-Frontal Circuits. Neuron, 63(5), 568–583. https://doi.org/10.1016/j.neuron.2009.08.028

Aru, J., Axmacher, N., Do Lam, A. T. A., Fell, J., Elger, C. E., Singer, W., & Melloni, L. (2012). Local Category-Specific Gamma Band Responses in the Visual Cortex Do Not Reflect Conscious Perception. Journal of Neuroscience, 32(43), 14909–14914. https://doi.org/10.1523/JNEUROSCI.2051-12.2012

Baars, B. J. (1998). Metaphors of consciousness and attention in the brain. Trends in Cognitive Sciences, 21, 58–62.

Block, N. (2011). Perceptual consciousness overflows cognitive access. Trends in cognitive sciences, 15(12), 567–575.

Block, N. (2019). What Is Wrong with the No-Report Paradigm and How to Fix It. Trends in Cognitive Sciences, 23(12), 1003–1013. https://doi.org/10.1016/j.tics.2019.10.001

Bogacz, R., Brown, E., Moehlis, J., Holmes, P., & Cohen, J. D. (2006). The physics of optimal decision making: A formal analysis of models of performance in two-alternative forced-choice tasks. Psychological Review, 113(4), 700–765. https://doi.org/10.1037/0033-295X.113.4.700

Boldt, A., & Yeung, N. (2015). Shared Neural Markers of Decision Confidence and Error Detection. Journal of Neuroscience, 35(8), 3478–3484. https://doi.org/10.1523/JNEUROSCI.0797-14.2015

Bollimunta, A., & Ditterich, J. (2012). Local Computation of Decision-Relevant Net Sensory Evidence in Parietal Cortex. Cerebral Cortex, 22(4), 903–917. https://doi.org/10.1093/cercor/bhr165

Brainard, D. H. (1997). The Psychophysics Toolbox. Spatial Vision, 10(4), 433–436. https://doi.org/10.1163/156856897X00357

Brown, R., Lau, H., & LeDoux, J. E. (2019). Understanding the Higher-Order Approach to Consciousness. Trends in Cognitive Sciences, 23(9), 754–768. https://doi.org/10.1016/j.tics.2019.06.009

Busemeyer, Jerome R. T., James T. (1993). Decision field theory: A dynamic-cognitive apporach to decision making in an uncertain environment. Psychological Review, 100(3), 432–459.

Chalmers, D. J. (1995). Facing Up to the Problem of Consciousness. Journal of Consciousness Studies, 2, 1–27.

Chaumon, M., Bishop, D. V. M., & Busch, N. A. (2015). A practical guide to the selection of independent components of the electroencephalogram for artifact correction. Journal of Neuroscience Methods, 250, 47–63. https://doi.org/10.1016/j.jneumeth.2015.02.025

Churchland, Anne. K., Kiani, R., Chaudhuri, R., Wang, X.-J., Pouget, A., & Shadlen, M. N. (2011). Variance as a Signature of Neural Computations during Decision Making. Neuron, 69(4), 818–831. https://doi.org/10.1016/j.neuron.2010.12.037

Dehaene, S., & Changeux, J.-P. (2011). Experimental and Theoretical Approaches to Conscious Processing. Neuron, 70(2), 200–227. https://doi.org/10.1016/j.neuron.2011.03.018

Dehaene, S., Changeux, J.-P., Naccache, L., Sackur, J., & Sergent, C. (2006). Conscious, preconscious, and subliminal processing: A testable taxonomy. Trends in Cognitive Sciences, 10(5), 204–211. https://doi.org/10.1016/j.tics.2006.03.007

Dehaene, S., Charles, L., King, J.-R., & Marti, S. (2014). Toward a computational theory of conscious processing. Current Opinion in Neurobiology, 25, 76–84. https://doi.org/10.1016/j.conb.2013.12.005

Delorme, A., & Makeig, S. (2004). EEGLAB: An open source toolbox for analysis of single-trial EEG dynamics including independent component analysis. Journal of Neuroscience Methods, 134(1), 9–21. https://doi.org/10.1016/j.jneumeth.2003.10.009

Devine, C. A., Gaffney, C., Loughnane, G. M., Kelly, S. P., & O’Connell, R. G. (2019). The role of premature evidence accumulation in making difficult perceptual decisions under temporal uncertainty. ELife, 8, e48526. https://doi.org/10.7554/eLife.48526

Ding, L., & Gold, J. I. (2010). Caudate Encodes Multiple Computations for Perceptual Decisions. Journal of Neuroscience, 30(47), 15747–15759. https://doi.org/10.1523/JNEUROSCI.2894-10.2010

Flavell, J. H. (1979). Metacognition and cognitive monitoring: A new area of cognitive-developmental inquiry. American Psychologist, 34(10), 906–911.

Fleming Stephen M., Dolan Raymond J. and Frith Christopher D. (2012) Metacognition: computation, biology and function. Phil. Trans. R. Soc. B 367: 1280–1286

Fleming, S. M., & Lau, H. C. (2014). How to measure metacognition. Frontiers in Human Neuroscience, 8. https://doi.org/10.3389/fnhum.2014.00443

Fleming, S. M., van der Putten, E. J., & Daw, N. D. (2018). Neural mediators of changes of mind about perceptual decisions. Nature Neuroscience, 21(4), 617–624. https://doi.org/10.1038/s41593-018-0104-6

Frassle, S., Sommer, J., Jansen, A., Naber, M., & Einhauser, W. (2014). Binocular Rivalry: Frontal Activity Relates to Introspection and Action But Not to Perception. Journal of Neuroscience, 34(5), 1738–1747. https://doi.org/10.1523/JNEUROSCI.4403-13.2014

Fu, Z., Wu, D.-A. J., Ross, I., Chung, J. M., Mamelak, A. N., Adolphs, R., & Rutishauser, U. (2019). Single-Neuron Correlates of Error Monitoring and Post-Error Adjustments in Human Medial Frontal Cortex. Neuron, 101(1), 165-177.e5. https://doi.org/10.1016/j.neuron.2018.11.016

Gherman, S., & Philiastides, M. G. (2015). Neural representations of confidence emerge from the process of decision formation during perceptual choices. NeuroImage, 106, 134–143. https://doi.org/10.1016/j.neuroimage.2014.11.036

Gold, J. I., & Shadlen, M. N. (2007). The Neural Basis of Decision Making. Annual Review of Neuroscience, 30(1), 535–574. https://doi.org/10.1146/annurev.neuro.29.051605.113038

Groppe, D. M., Bickel, S., Dykstra, A. R., Wang, X., Mégevand, P., Mercier, M. R., Lado, F. A., Mehta, A. D., & Honey, C. J. (2017). iELVis: An open source MATLAB toolbox for localizing and visualizing human intracranial electrode data. Journal of Neuroscience Methods, 281, 40–48. https://doi.org/10.1016/j.jneumeth.2017.01.022

Hanks, T. D., Kopec, C. D., Brunton, B. W., Duan, C. A., Erlich, J. C., & Brody, C. D. (2015). Distinct relationships of parietal and prefrontal cortices to evidence accumulation. Nature, 520(7546), 220–223. https://doi.org/10.1038/nature14066

Herding, J., Ludwig, S., von Lautz, A., Spitzer, B., & Blankenburg, F. (2019). Centro-parietal EEG potentials index subjective evidence and confidence during perceptual decision making. NeuroImage, 201, 116011. https://doi.org/10.1016/j.neuroimage.2019.116011

Kaminski, J., Sullivan, S., Chung, J. M., Ross, I. B., Mamelak, A. N., & Rutishauser, U. (2017). Persistently active neurons in human medial frontal and medial temporal lobe support working memory. Nature Neuroscience, 20(4), 590–601. https://doi.org/10.1038/nn.4509

Kang, Y. H. R., Petzschner, F. H., Wolpert, D. M., & Shadlen, M. N. (2017). Piercing of Consciousness as a Threshold-Crossing Operation. Current Biology, 27(15), 2285-2295.e6. https://doi.org/10.1016/j.cub.2017.06.047

Katz, L. N., Yates, J. L., Pillow, J. W., & Huk, A. C. (2016). Dissociated functional significance of decision-related activity in the primate dorsal stream. Nature, 535(7611), 285–288. https://doi.org/10.1038/nature18617

Kiani, R., & Shadlen, M. N. (2009). Representation of Confidence Associated with a Decision by Neurons in the Parietal Cortex. Science, 324(5928), 759–764. https://doi.org/10.1126/science.1169405

Kleiner, M. (n.d.). What’s new in Psychtoolbox-3? 89.

Koch, C., Massimini, M., Boly, M., & Tononi, G. (2016). Neural correlates of consciousness: Progress and problems. Nature Reviews Neuroscience, 17(5), 307–321. https://doi.org/10.1038/nrn.2016.22

Koriat, A. (2006). Metacognition and consciousness. In The Cambridge Handbook of Consciousness (Vol. 3, pp. 289–326).

Kvam, P. D., Pleskac, T. J., Yu, S., & Busemeyer, J. R. (2015). Interference effects of choice on confidence: Quantum characteristics of evidence accumulation. Proceedings of the National Academy of Sciences, 112(34), 10645–10650. https://doi.org/10.1073/pnas.1500688112

Lamme, V. A. F. (2010). How neuroscience will change our view on consciousness. Cognitive Neuroscience, 1(3), 204–220. https://doi.org/10.1080/17588921003731586

Lau, H., & Rosenthal, D. (2011). Empirical support for higher-order theories of conscious awareness. Trends in Cognitive Sciences, 15(8), 365–373. https://doi.org/10.1016/j.tics.2011.05.009

Li, Q., Hill, Z., & He, B. J. (2014). Spatiotemporal Dissociation of Brain Activity Underlying Subjective Awareness, Objective Performance and Confidence. Journal of Neuroscience, 34(12), 4382–4395. https://doi.org/10.1523/JNEUROSCI.1820-13.2014

Makeig, S., Bell, A. J., Jung, T.-P., & Sejnowski, T. J. (n.d.). Independent Component Analysis of Electroencephalographic Data. 7.

Maniscalco, B., & Lau, H. (2016). The signal processing architecture underlying subjective reports of sensory awareness. Neuroscience of Consciousness, 2016(1). https://doi.org/10.1093/nc/niw002

Mashour, G. A., Roelfsema, P., Changeux, J. P., & Dehaene, S. (2020). Conscious processing and the global neuronal workspace hypothesis. Neuron, 105(5), 776–798.

Mazor, M., Friston, K. J., & Fleming, S. M. (2020). Distinct neural contributions to metacognition for detecting, but not discriminating visual stimuli. ELife, 9, e53900. https://doi.org/10.7554/eLife.53900

Meuwese, J. D. I., van Loon, A. M., Lamme, V. A. F., & Fahrenfort, J. J. (2014). The subjective experience of object recognition: Comparing metacognition for object detection and object categorization. Attention, Perception, & Psychophysics. https://doi.org/10.3758/s13414-014-0643-1

Moutard, C., Dehaene, S., & Malach, R. (2015). Spontaneous Fluctuations and Non-linear Ignitions: Two Dynamic Faces of Cortical Recurrent Loops. Neuron, 88(1), 194–206. https://doi.org/10.1016/j.neuron.2015.09.018

Nagel, T. (1974). What is it like to be a bat. The Philosophical Review, 4, 435–450.

O’Connell, R. G., Dockree, P. M., & Kelly, S. P. (2012). A supramodal accumulation-to-bound signal that determines perceptual decisions in humans. Nature Neuroscience, 15(12), 1729–1735. https://doi.org/10.1038/nn.3248

Odegaard, B., Grimaldi, P., Cho, S. H., Peters, M. A. K., Lau, H., & Basso, M. A. (2018). Superior colliculus neuronal ensemble activity signals optimal rather than subjective confidence. Proceedings of the National Academy of Sciences, 115(7), E1588–E1597. https://doi.org/10.1073/pnas.1711628115

Pearson, R., & Powell, T. (1985). The projection of the primary somatic sensory cortex upon area 5 in the monkey. Brain Research Reviews, 9(1), 89–107. https://doi.org/10.1016/0165-0173(85)90020-7

Pelli, D. G. (1997). The VideoToolbox software for visual psychophysics: Transforming numbers into movies. Spatial Vision, 10(4), 437–442.

Pereira, M., Faivre, N., Iturrate, I., Wirthlin, M., Serafini, L., Martin, S., Desvachez, A., Blanke, O., Van De Ville, D., & Millán, J. del R. (2020). Disentangling the origins of confidence in speeded perceptual judgments through multimodal imaging. Proceedings of the National Academy of Sciences, 117(15), 8382–8390. https://doi.org/10.1073/pnas.1918335117

Philiastides, M. G., Heekeren, H. R., & Sajda, P. (2014). Human Scalp Potentials Reflect a Mixture of Decision-Related Signals during Perceptual Choices. Journal of Neuroscience, 34(50), 16877–16889. https://doi.org/10.1523/JNEUROSCI.3012-14.2014

Phillips, I., & Morales, J. (2020). The Fundamental Problem with No-Cognition Paradigms. Trends in Cognitive Sciences, 24(3), 165–167. https://doi.org/10.1016/j.tics.2019.11.010

Pitts, M. A., Padwal, J., Fennelly, D., Martínez, A., & Hillyard, S. A. (2014). Gamma band activity and the P3 reflect post-perceptual processes, not visual awareness. NeuroImage, 101, 337–350. https://doi.org/10.1016/j.neuroimage.2014.07.024

Pleskac, T. J., & Busemeyer, J. R. (2010). Two-stage dynamic signal detection: A theory of choice, decision time, and confidence. Psychological Review, 117(3), 864–901. https://doi.org/10.1037/a0019737

Quiroga, R. Q., Mukamel, R., Isham, E. A., Malach, R., & Fried, I. (2008). Human single-neuron responses at the threshold of conscious recognition. Proceedings of the National Academy of Sciences, 105(9), 3599–3604. https://doi.org/10.1073/pnas.0707043105

Ratcliff, R., & Van Dongen, H. P. A. (2011). Diffusion model for one-choice reaction-time tasks and the cognitive effects of sleep deprivation. Proceedings of the National Academy of Sciences, 108(27), 11285–11290. https://doi.org/10.1073/pnas.1100483108

Ratcliff, Roger, & Rouder, J. N. (n.d.). A Diffusion Model Account of Masking in Two-Choice Letter Identification. 14.

Reber, T. P., Faber, J., Niediek, J., Boström, J., Elger, C. E., & Mormann, F. (2017). Single-Neuron Correlates of Conscious Perception in the Human Medial Temporal Lobe. Current Biology, 27(19), 2991-2998.e2. https://doi.org/10.1016/j.cub.2017.08.025

Roitman, J. D., & Shadlen, M. N. (2002). Response of Neurons in the Lateral Intraparietal Area during a Combined Visual Discrimination Reaction Time Task. The Journal of Neuroscience, 22(21), 9475–9489. https://doi.org/10.1523/JNEUROSCI.22-21-09475.2002

Rutishauser, U., Aflalo, T., Rosario, E. R., Pouratian, N., & Andersen, R. A. (2018). Single-Neuron Representation of Memory Strength and Recognition Confidence in Left Human Posterior Parietal Cortex. Neuron, 97(1), 209-220.e3. https://doi.org/10.1016/j.neuron.2017.11.029

Rutishauser, U., Schuman, E. M., & Mamelak, A. N. (2006). Online detection and sorting of extracellularly recorded action potentials in human medial temporal lobe recordings, in vivo. Journal of Neuroscience Methods, 154(1–2), 204–224. https://doi.org/10.1016/j.jneumeth.2005.12.033

Rutishauser, U., Ye, S., Koroma, M., Tudusciuc, O., Ross, I. B., Chung, J. M., & Mamelak, A. N. (2015). Representation of retrieval confidence by single neurons in the human medial temporal lobe. Nature Neuroscience, 18(7), 1041–1050. https://doi.org/10.1038/nn.4041

Salti, M., Monto, S., Charles, L., King, J.-R., Parkkonen, L., & Dehaene, S. (2015). Distinct cortical codes and temporal dynamics for conscious and unconscious percepts. ELife, 4, e05652. https://doi.org/10.7554/eLife.05652

Savitzky, Abraham., & Golay, M. J. E. (1964). Smoothing and Differentiation of Data by Simplified Least Squares Procedures. Analytical Chemistry, 36(8), 1627–1639. https://doi.org/10.1021/ac60214a047

Shea, N., & Frith, C. D. (2019). The Global Workspace Needs Metacognition. Trends in Cognitive Sciences, 23(7), 560–571. https://doi.org/10.1016/j.tics.2019.04.007

Tagliabue, C. F., Veniero, D., Benwell, C. S. Y., Cecere, R., Savazzi, S., & Thut, G. (2019). The EEG signature of sensory evidence accumulation during decision formation closely tracks subjective perceptual experience. Scientific Reports, 9(1), 4949. https://doi.org/10.1038/s41598-019-41024-4

Tsuchiya, N., Wilke, M., Frässle, S., & Lamme, V. A. F. (2015). No-Report Paradigms: Extracting the True Neural Correlates of Consciousness. Trends in Cognitive Sciences, 19(12), 757–770. https://doi.org/10.1016/j.tics.2015.10.002

van den Berg, R., Anandalingam, K., Zylberberg, A., Kiani, R., Shadlen, M. N., & Wolpert, D. M. (2016). A common mechanism underlies changes of mind about decisions and confidence. ELife, 5, e12192. https://doi.org/10.7554/eLife.12192

van Vugt, B., Dagnino, B., Vartak, D., Safaai, H., Panzeri, S., Dehaene, S., & Roelfsema, P. R. (2018). The threshold for conscious report: Signal loss and response bias in visual and frontal cortex. Science, 360(6388), 537–542. https://doi.org/10.1126/science.aar7186

Wyart, V., & Tallon-Baudry, C. (2009). How Ongoing Fluctuations in Human Visual Cortex Predict Perceptual Awareness: Baseline Shift versus Decision Bias. Journal of Neuroscience, 29(27), 8715–8725. https://doi.org/10.1523/JNEUROSCI.0962-09.2009

Zeki, S. (2007). The disunity of consciousness. In Progress in Brain Research (Vol. 168, pp. 11–268). Elsevier. https://doi.org/10.1016/S0079-6123(07)68002-9

Zhou, Y., & Freedman, D. J. (2019). Posterior parietal cortex plays a causal role in perceptual and categorical decisions. Science, 180–185.

