## Supplementary results for "Evidence accumulation determines conscious access"

**Supplementary behavioral results**

*Experiments 1 - 2*

We verified that stimulus intensity did not vary significantly between hits and misses during Experiment 1 (t(272)=0.52, p=0.60) nor during Experiment 2 (t(101)=1.36, p=0.18). In Experiment 2, a response was considered correct when the participant answered “yes” following a stimulus (hit) or “no” in the absence of a stimulus (correct rejection): hits and correct rejections represented respectively 39.69% and 20.61% of trials. Incorrect trials included “no” responses following a stimulus (miss) and “yes” responses in the absence of a stimulus (false alarms), which represented respectively 38.93% and 0.80%. This corresponded to a d-prime of 1.64 and a bias of 1.64 indicative of conservative behavior (i.e., tendency to answer no in the presence of a stimulus) and consistent with healthy controls described below. Importantly, stimulus onset did not vary significantly between hits and misses (t(101)=-0.59, p=0.56). Confidence was analyzed separately for “yes” and “no” responses.

*Experiment 4*

A response was considered correct when participants answered “yes” following a stimulus (hit) or “no” in the absence of a stimulus (correct rejection): hits and correct rejections represented respectively 37.81%±1.70 and 18.9%±0.66 of trials. Incorrect trials included “no” responses following a stimulus (miss) and “yes” responses in the absence of a stimulus (false alarms), which represented respectively 42.07%±1.67 and 1.37%±0.71 of trials. This corresponded to an average d-prime of 1.67±0.26, and a bias of 7.62±3.53 indicative of conservative behavior (i.e., tendency to answer no in the presence of a stimulus). Importantly, stimulus intensity and stimulus onset did not vary significantly between hits and misses (t(17)=0.81, p=0.43 and t(17)=0.12, p=0.91 , respectively). Confidence was analyzed separately for “yes” and “no” responses. For “yes” responses, average confidence was higher in hits (77.79%±3.82) than in false alarms (62.16%±7.15; $t(23.12)=-3.78$, $p=0.001$). For “no” responses, confidence was higher in correct rejections (86.31%±3.47) than in misses (83.06%±3.44; $t(17)=5.59$, $p<0.001$). These results indicate higher confidence for correct than incorrect responses (confidence gap: t(32)=4.10, p<0.001), implying that participants had a metacognitive access to tactile detection processes.

**Supplementary electrophysiological results**

*Analyses on population of selective neurons for Experiments 1–3*

We sought to verify the consistency of neuronal responses for selective neurons. We first analyzed averaged normalized responses for each experiment. For each selective neuron with an increase in firing rate for hits (N=47 for Experiment 1, N=10 for Experiment 2) or for increased amplitudes (N=14 for Experiment 3), we divided firing rates by the firing rate during the 300 ms pre-stimulus baseline, averaged across trials. This scaling of the firing rates allowed for better comparisons in neurons with different firing rates. We found similar responses as described single neurons (Fig. S2A,B,C).

*Choice probabilities (Experiment 2)*

We confirmed the factorial analysis reported in the main text by probing whether the specificity for detection correlated with that of confidence using neuronal choice probabilities (CP). Choice probability was computed using the area under the curve (AUC) for either detection or confidence and normalized to the [-1,1] interval: CP = 2*AUC-1. A significant correlation was found between CP for detection and confidence among selective neurons for hits (R=0.66, p=0.0052), but not for misses (R=0.27, p=0.28).

**Supplementary modelling results**

*Analysis of confidence variance (Experiment 1)*

To further support our evidence accumulation model, we verified some of its predictions a-posteriori. Notably, our model predicted lower variance for simulated confidence in misses (0.012±0.001) than in hits (0.026±0.002; z=3.2, p=0.0012), since accumulated evidence was bounded between zero and the decision bound for misses but not for hits. Similarly, in the data, confidence ratings had lower variance for misses (0.015±0.002) compared to hits (0.026±0.002; z=2.6, p=0.0084). Furthermore, since the drift rate was null for correct rejections, the model predicted even lower variance of confidence for correct rejections (0.007±0.001) compared to misses (z=3.7; p=0.002), which we confirmed in the data: (0.011±0.001 vs. 0.015±0.002; z=3.1; p=0.0018).

The predictions of our maximal evidence model correlated with metacognitive sensitivity observed in the data for both “yes” responses (R=0.60, p=0.001, permutation test) and “no” responses (R=0.61, p=0.009; Fig.S8b), showing that our model successfully predicted metacognitive performance.

Interestingly, evidence accumulation models such as ours could naturally explain suboptimal confidence ratings or metacognitive noise (Maniscalco & Lau, 2016) through the influence of first-order parameters on post-decisional evidence accumulation used for confidence readouts. In our model in particular, for “yes” answers, confidence is sampled at a variable timing after the decision. This timing depends on a combination of first-order parameters, including leakage. According to the model, accumulation processes characterized by strong leakage have sharper evidence maxima, leading to a sampling of evidence close in time to the decision and therefore, low metacognitive noise. On the contrary, accumulation processes with hardly no leakage result in evidence levels that oscillate randomly around the level reached after no more evidence is available and thus have a higher metacognitive noise. We verified this hypothesis a-posteriori: the leakage parameter correlated with the metacognitive sensitivity (AUC) for “yes” responses (R=0.50, p=0.029).

### Supplementary figures

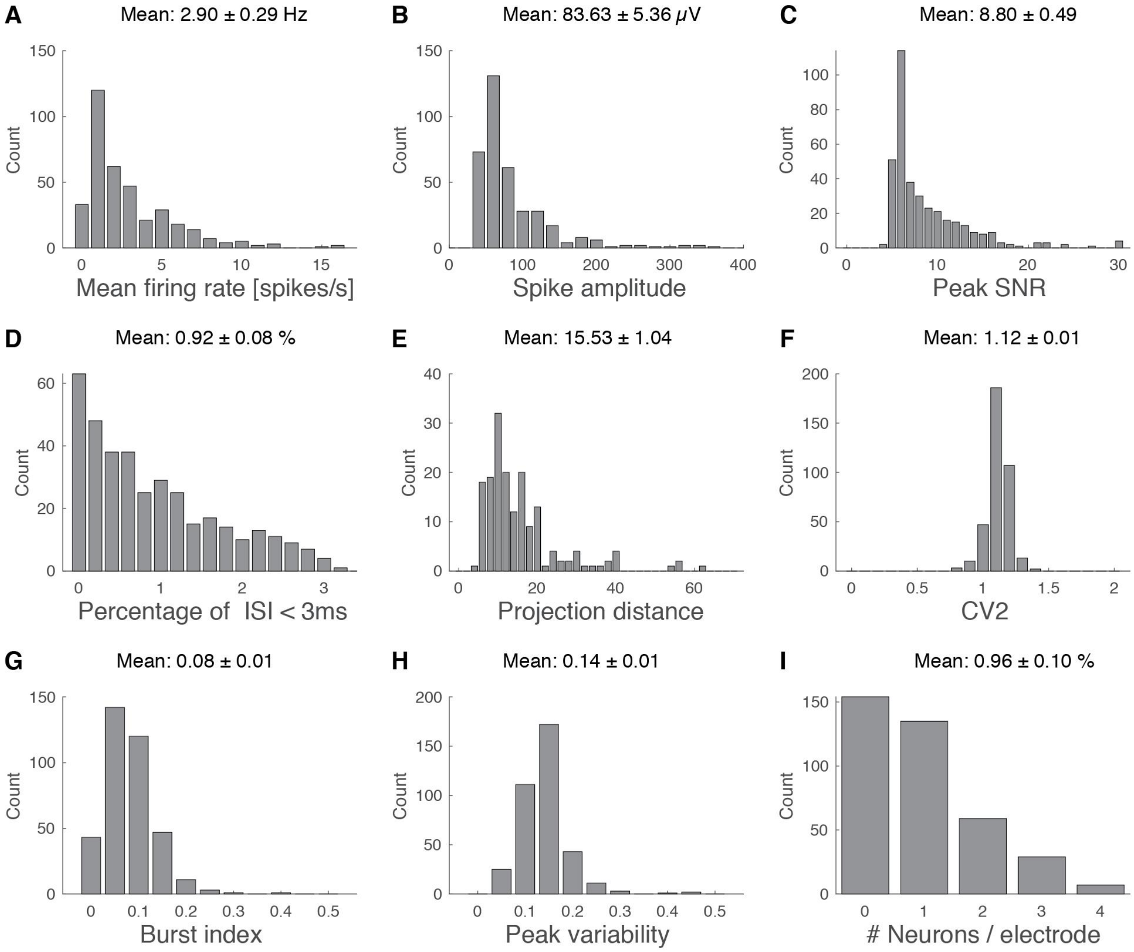


***Fig. S1.*** *Spike sorting statistics.* ***(A)*** *Histogram of firing rates.* ***(B)*** *Histogram of mean spike amplitudes (at the peak, rectified, since all neurons had negative action potentials).* ***(C)*** *Histogram of signal-to-noise ratio (ratio between spike peak amplitude and estimated noise).* ***(D)*** *Histogram of the percentage of inter-spike interval lower than a 3 ms refractory period.* ***(E)*** *Histogram of projection distance (Rutishauser et al., 2006) for electrodes with more than one single unit. (****F****) Histogram of modified coefficient of variation.* ***(G)*** *Histogram of burst index (number of inter-spike intervals lower than 10 ms).* ***(H)*** *Histogram of peak variability (ratio between the standard deviation of spike amplitude and the mean spike amplitude).* ***(I)*** *Histogram of number of neurons found per electrode.*

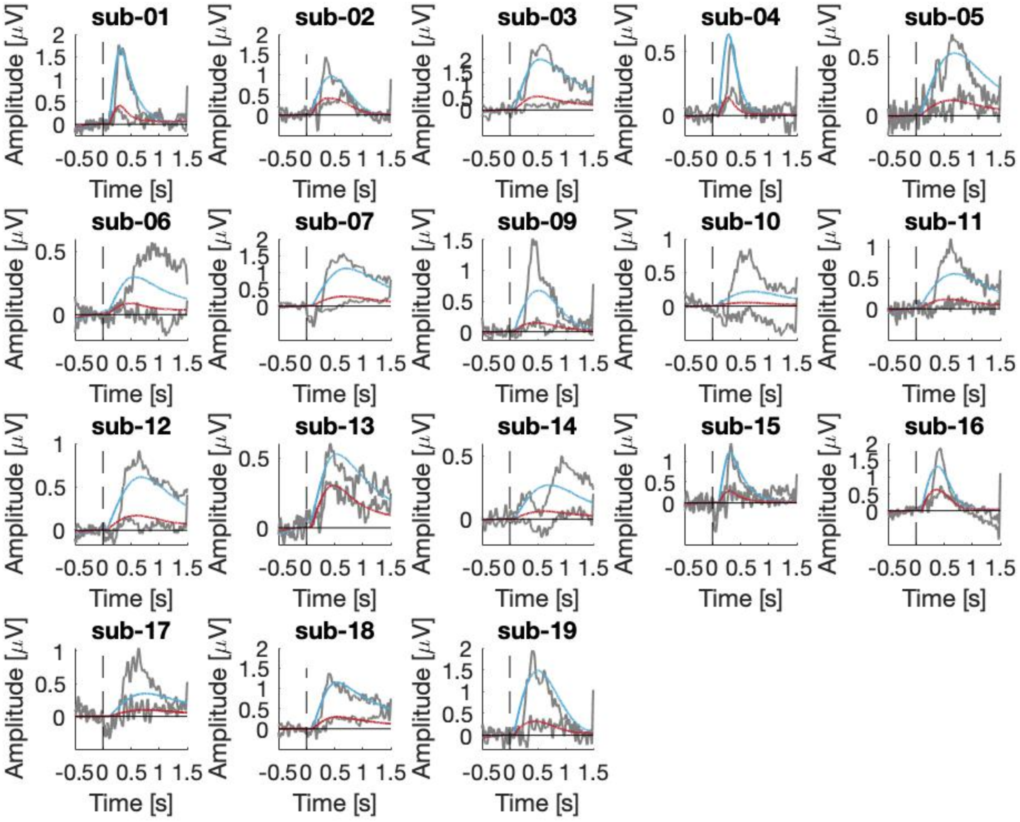


***Fig S2.*** *Model fits in Experiment 4. Each plot represents the individual fit of EEG data for hits (cyan traces) and misses (red traces). Observed data are shown in grey. Vertical dashed line represents stimulus onset. Average R across participants: 0.70±0.03.*

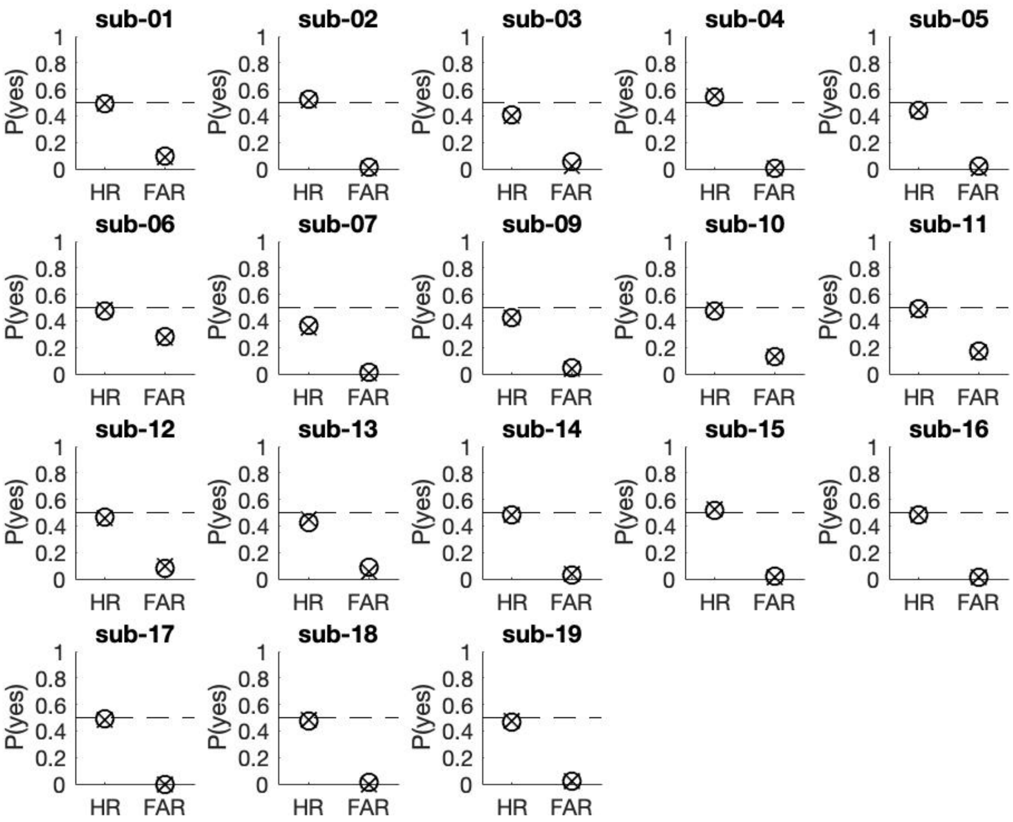


***Fig S3.*** *Model fits in Experiment 4. Each plot represents individual fits of detection reports, summarized as hit rate (HR) and false alarm rate (FAR). Circles show data and crosses show model simulations. Horizontal dashed shows the 50% hit rate target. Correlation: R between participants: 0.96 for hit rate and 0.95 for false alarm rate.*

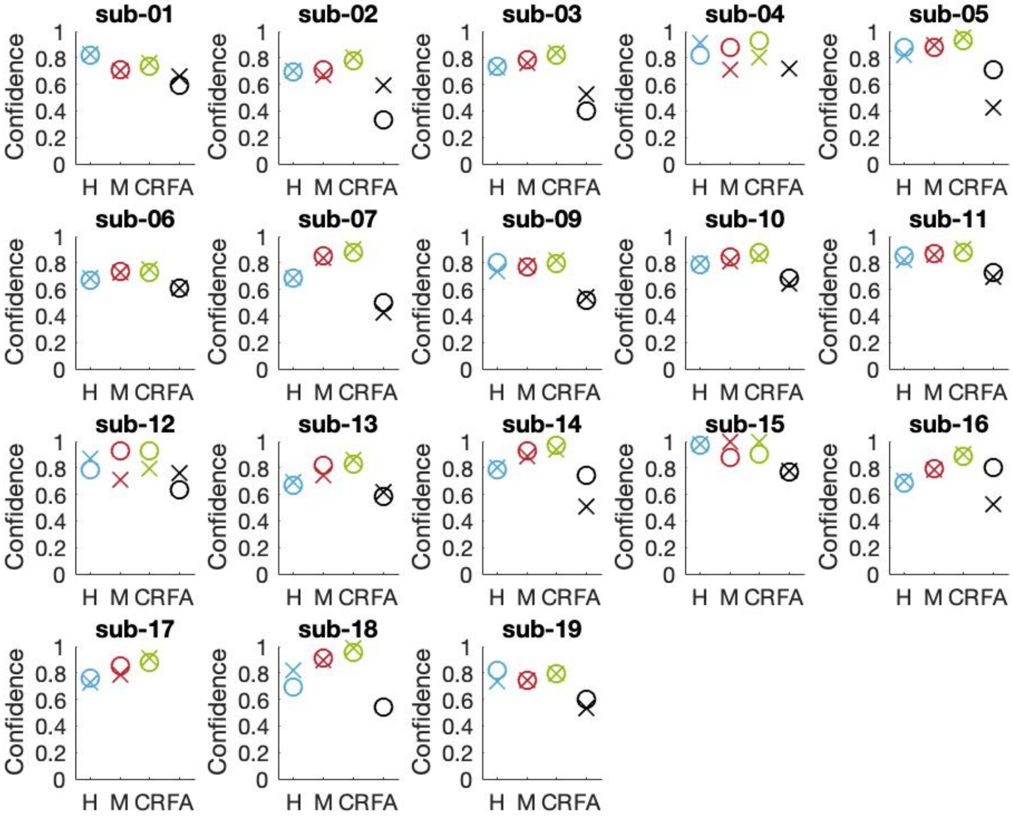


***Fig S4.*** *Model fits in Experiment 4. Each plot represents individual fits of confidence ratings for hits (H), misses (M), correct rejections (CR) and false alarms (FA; when any). Circles show data and crosses show model simulations. Average R across participants 0.83±0.03 for hits, 0.85±0.03 for misses, 0.81±0.04 for correct rejections and 0.45±0.09 for false alarms.*

**Supplementary references**

Maniscalco, B., & Lau, H. (2016). The signal processing architecture underlying subjective reports of sensory awareness. *Neuroscience of Consciousness*, *2016*(1). https://doi.org/10.1093/nc/niw002

Rutishauser, U., Schuman, E. M., & Mamelak, A. N. (2006). Online detection and sorting of extracellularly recorded action potentials in human medial temporal lobe recordings, in vivo. *Journal of Neuroscience Methods*, *154*(1–2), 204–224. https://doi.org/10.1016/j.jneumeth.2005.12.033
